# An inactivation switch enables rhythms in a *Neurospora* clock model

**DOI:** 10.1101/599506

**Authors:** Abhishek Upadhyay, Michael Brunner, Hanspeter Herzel

**Affiliations:** Institute for Theoretical Biology, Charité – Universitätsmedizin Berlin and Humboldt University of Berlin, Philippstr. 13, 10115, Berlin, Germany; Biochemistry Center, University of Heidelberg, Im Neuenheimer Feld 328, 69120, Heidelberg, Germany

**Keywords:** Circadian clock, Mathematical modelling, Molecular switch, *Neurospora crassa*, Glucose compensation

## Abstract

An autonomous endogenous time-keeping is ubiquitous across many living organisms known as circadian clock when it has a period of about 24 hours. Interestingly, the fundamental design principle with a network of interconnected negative and positive feedback loops is conserved through evolution, although the molecular components differ. Filamentous fungus *Neurospora crassa* is a well established chrono-genetics model organism to investigate the underlying mechanisms. The core negative feedback loop of the clock of *Neurospora* is composed of the transcription activator White Collar Complex (WCC) (heterodimer of WC1 and WC2) and the inhibitory element called FFC complex which is made of FRQ (Frequency protein), FRH (Frequency interacting RNA Helicase) and CK1a (Casein kinase 1a). While exploring their temporal dynamics we investigate how limit cycle oscillations arise and how molecular switches support self-sustained rhythms. We develop a mathematical model of 10 variables with 26 parameters to understand the interactions and feedbacks among WC1 and FFC elements in nuclear and cytoplasmic compartments. We performed control and bifurcation analysis to show that our novel model produces robust oscillations with a wild-type period of 22.5 hrs. Our model reveals a switch between WC1 induced transcription and FFC assisted inactivation of WC1. Using the new model we also study possible mechanisms of glucose compensation. A fairly simple model with just 3 non-linearities helps to elucidate clock dynamics revealing a mechanism of rhythms production. The model can further be utilized to study entrainment and temperature compensation.

## 1. Introduction

The earth’s rotation around its own axis gives rise to the 24 hours day and night cycles. To anticipate these daily environmental changes in light and temperature, life in all its kingdoms has evolved a time keeping molecular machinery [1–3]. In humans, disruption in the clock functioning due to jet lag, shift work, light at night or other reasons increases the risk of diseases such as cancer, obesity and brain disorders [4].

In order to understand basic clock mechanisms we study a filamentous fungus, *Neurospora crassa* (*N. crassa*) [5]. Their natural habitats are soils, plants, trees and food resources [6,7]. The ability of *N. crassa* to deconstruct and metabolize plant cell walls is crucial for environmental carbon and other nutrient cycling. There are also various potential biotechnological applications such as the production of biofuels. An example is the production of ethanol from xylose derived from plant dry matter biomass (lignocellulosic substrates) [8]. The xylan to ethanol pathway has been found to be clock regulated at each stage [9]. Mathematical modelling is useful to understand the *Neurospora* clock mechanism with multiple known feedbacks [10–13].

Figure 1 demonstrates the transcription-translation feedback loop which consists of the inhibitory protein Frequency (FRQ), the activating transcription factor White Collar Complex (WCC), the FRQ stabilizer Frequency-interacting Helicase (FRH), and the Casein Kinase-1a (CK1a). Oscillations are based on delayed negative feedbacks. In *Neurospora*, clock gene frequency inhibits its own transcription after intermediate steps such as transcription, translation, dimerization, phosphorylations and nuclear import. Moreover, stabilization of FRQ is governed by FRH and CK1a binding in forming the FFC complex. Later, FFC is involved in the inhibition of WCC induced transcription [14–18].

**Figure 1.**
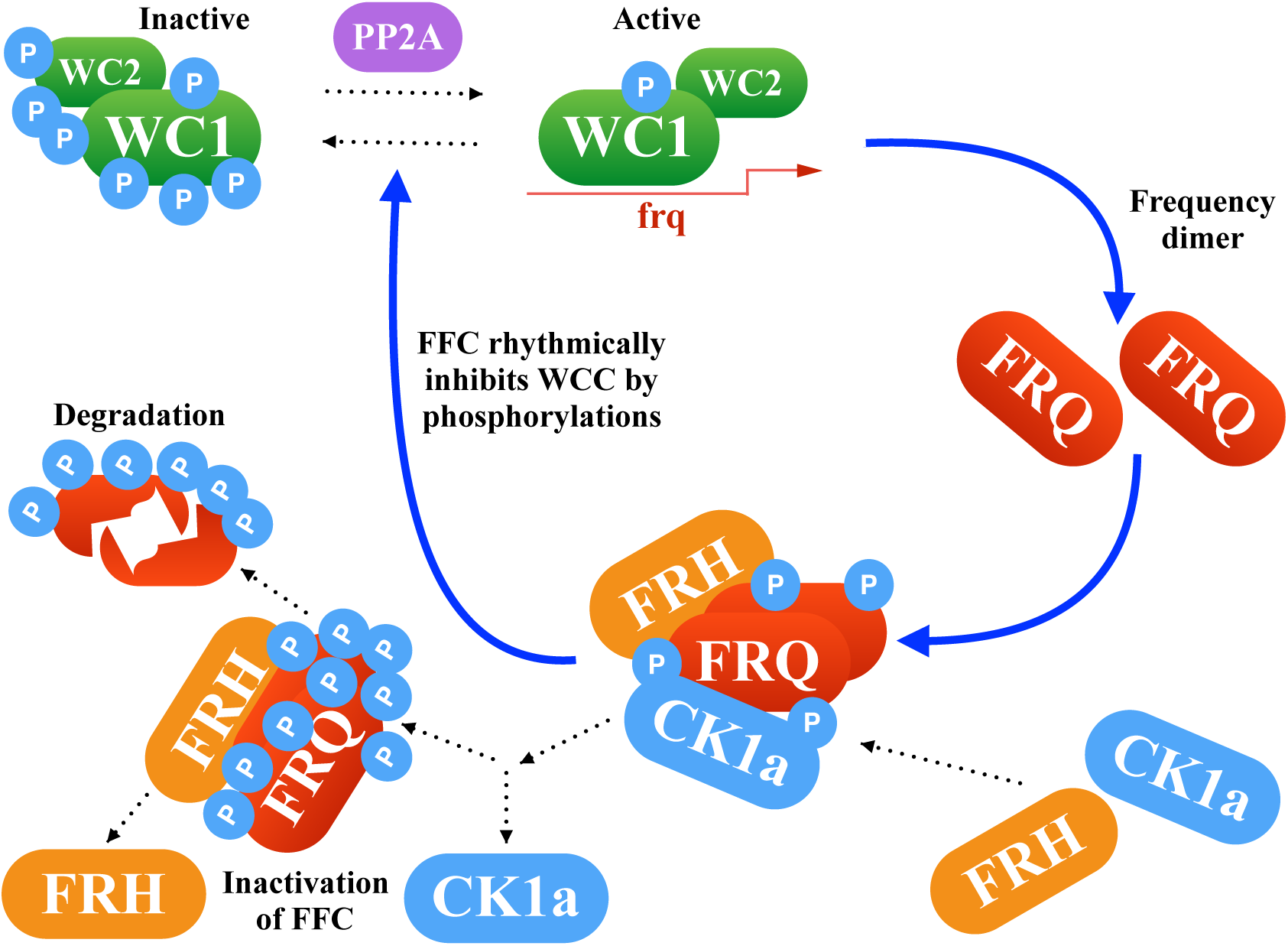
Core network of *Neurospora* clock: The delayed negative feedback via FRQ is controlled by complex formations and phosphorylations.

The detailed mechanisms of the negative feedback are not well understood. For example, the inhibition of WCC via FFC might involve hyperphosphorylation, sequestration and stoichoimetric inhibition [19–24]. Moreover, it is not clear how the long delay is realized to obtain daily rhythms. In order to obtain self-sustained oscillations nonlinearities are necessary. We explore the role of nonlinearities in the system and try to identify switch mechanisms. Finally, we discuss possible functions of the documented positive feedback in connection with glucose compensation. A comprehensive analysis of our mathematical model reveals that an inactivation switch allows self sustained rhythms in the *Neurospora* clock. Our model also suggests in-built glucose compensation mechanisms.

## 2. Results

### 2.1. Modelling the core clock elements

The negative feedback loop involves an implicit delay consisting of frq transcription and translation, FRQ dimerization, CK1a binding, nuclear import, formation of the FFC complex, and FFCn interaction with WC1n. A positive feedback involves the nuclear export of FFC supporting WC1c translation [25,26].

It is well known that FRQ and WCC are phosphorylated at many sites. Rhythmic phosphorylation influences complex formation and stability [27,28]. In our current model we take these phosphorylations implicitly into account via appropriate rate constants.

Figure 2 shows the turnover of frq, wcc and the associated negative and positive feedbacks. Using primarily linear kinetic terms this scheme can be directly translated into a system of ordinary differential equations (ODEs) [29–31]. These equations are presented in Appendix A1. The 10 time-varying concentrations are denoted with brackets and 26 kinetic parameters are enumerated as a01, a02, a1, a2, …, a21, n, A, A2.

**Figure 2.**
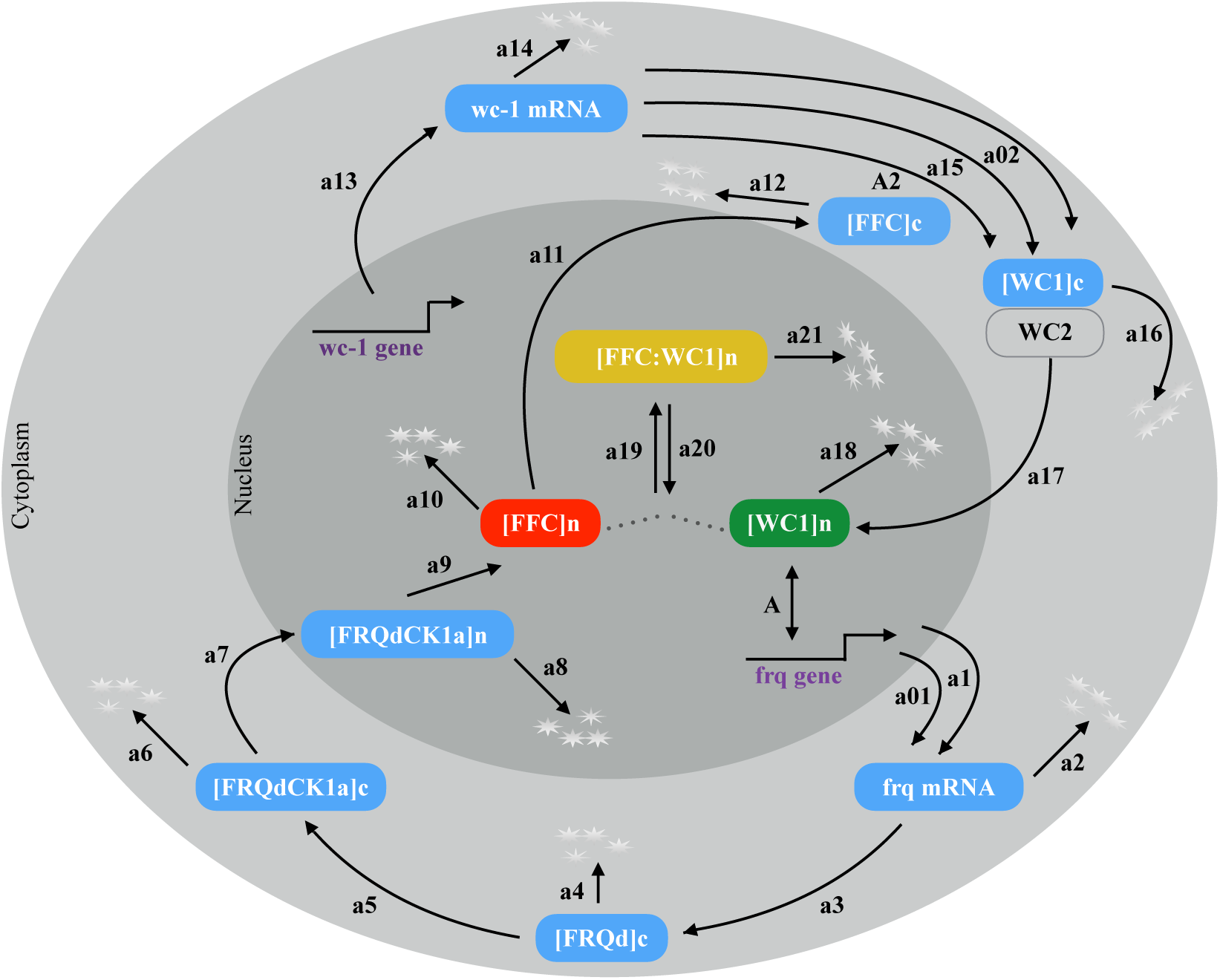
*Neurospora* circadian clock model: Wiring diagram of the model shows compartmentalization into nucleus and cytoplasm, turnover of frq and wc1, complex formations and nuclear translocation.

### 2.2. The model reproduces self sustained rhythms

The basic scheme of the model is adapted from a previous study [32]. In particular, we kept 22 parameters as in the original publication. In order to study the role of FRH and FFC, we introduced additional variables. The corresponding degradation rates were taken from experiments [27,33]. The remaining 2 unknown parameters were adjusted to get a period of 22.5 hours and robust limit cycle oscillations (see Appendix A2).

Our resulting model consists of 10 ordinary differential equations (ODEs) and 26 kinetic parameters. In order to get self-sustained rhythms nonlinearities are necessary. Inspection of the equations in ‘Appendix B: A1’ reveals that there are just 3 nonlinear terms: The transcription of the gene frq is modeled via a quadratic Hill function representing multiple WCC binding sites. The complex formation of FFC and WC1 is described by a bilinear term. The third nonlinearity models the positive feedback of the FFCc complex on WC1c translation as in the original model [32].

Figure 3 shows simulations of our model for the default parameters listed in Appendix A2. The oscillations of Frequency mRNA and protein (blue) are almost sinusoidal whereas the levels of the inhibitory complex FFCn (red) and the transcription factor WC1n (green) display spike-like behaviour. The resulting FFCnWC1n complex (yellow) exhibits even two peaks during one circadian cycle. This observation illustrates that nonlinearities can generate pronounced “harmonics”, an example of ultradian rhythms. Harmonics have been described in large-scale transcription profiles in mouse [34,35] and recently also in *Neurospora* [36]. The final graph in Figure 3 represents the transcription of the frq gene. It is evident that there is a sharp ON/OFF switch of transcription. In a later section we relate this temporal switch to FFC-assisted inactivation of the transcription factor WC1.

**Figure 3.**
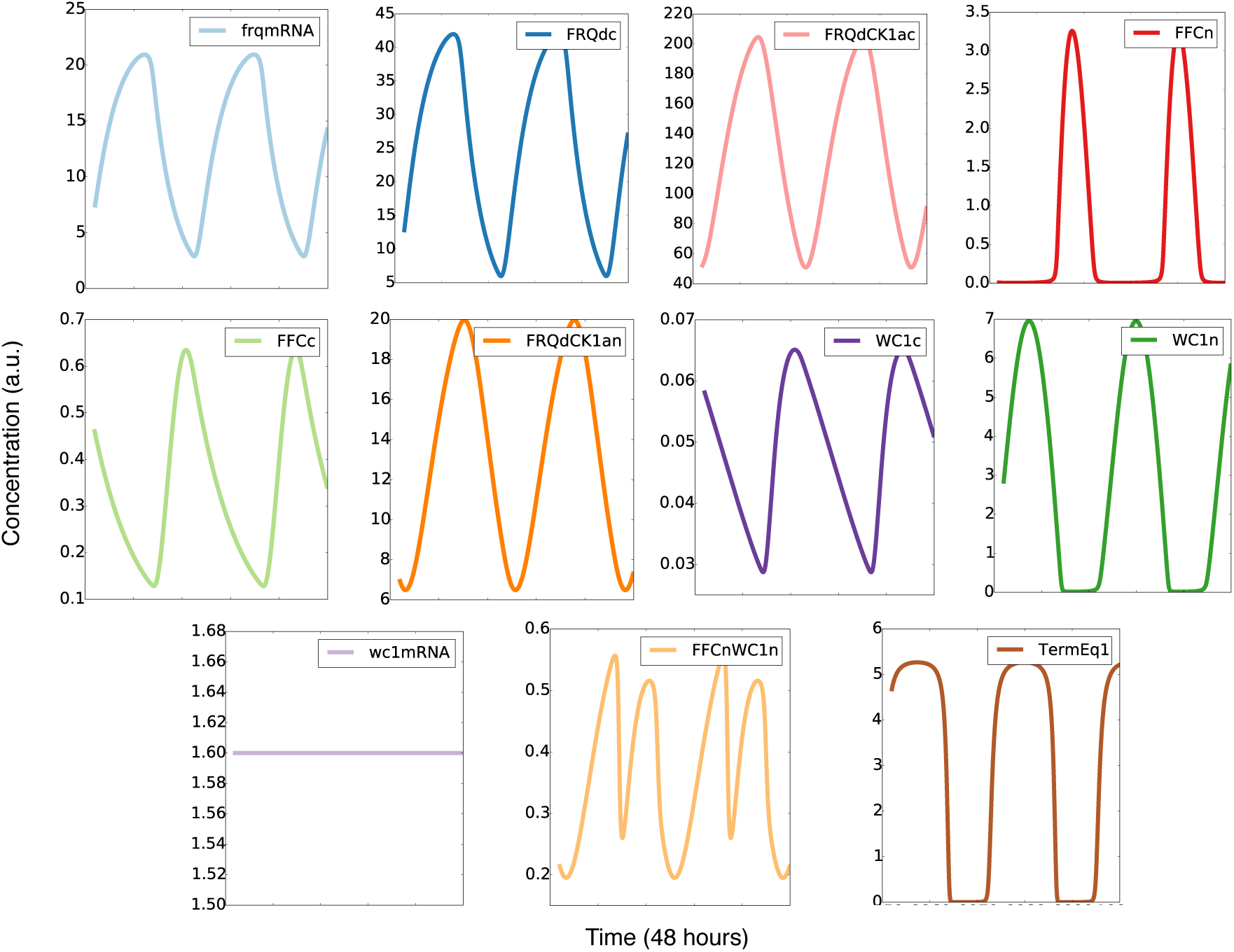
Simulated time series: There are sinusoidal and spike-like waveforms, harmonics and a temporal switch (see text).

### 2.3. Bifurcation analysis of our model

The available kinetic data are not sufficient to determine all model parameters precisely. Moreover, external conditions can induce parameter variations. Consequently, we varied all 26 parameters in a comprehensive manner to explore the robustness of the model and to evaluate the role of the different kinetic terms along the lines of previous studies [37,38].

It turns out that self-sustained oscillations are obtained in wide ranges of parameters (see Figure 4 and Appendix A3). Amplitudes and periods vary smoothly with most parameters. The onset of oscillations occurs in most cases via a Hopf bifurcation, a transition from damped to self-sustained rhythms. Close to Hopf bifurcations we find large amplitude variations but weaker dependencies of the period on parameters.

**Figure 4.**
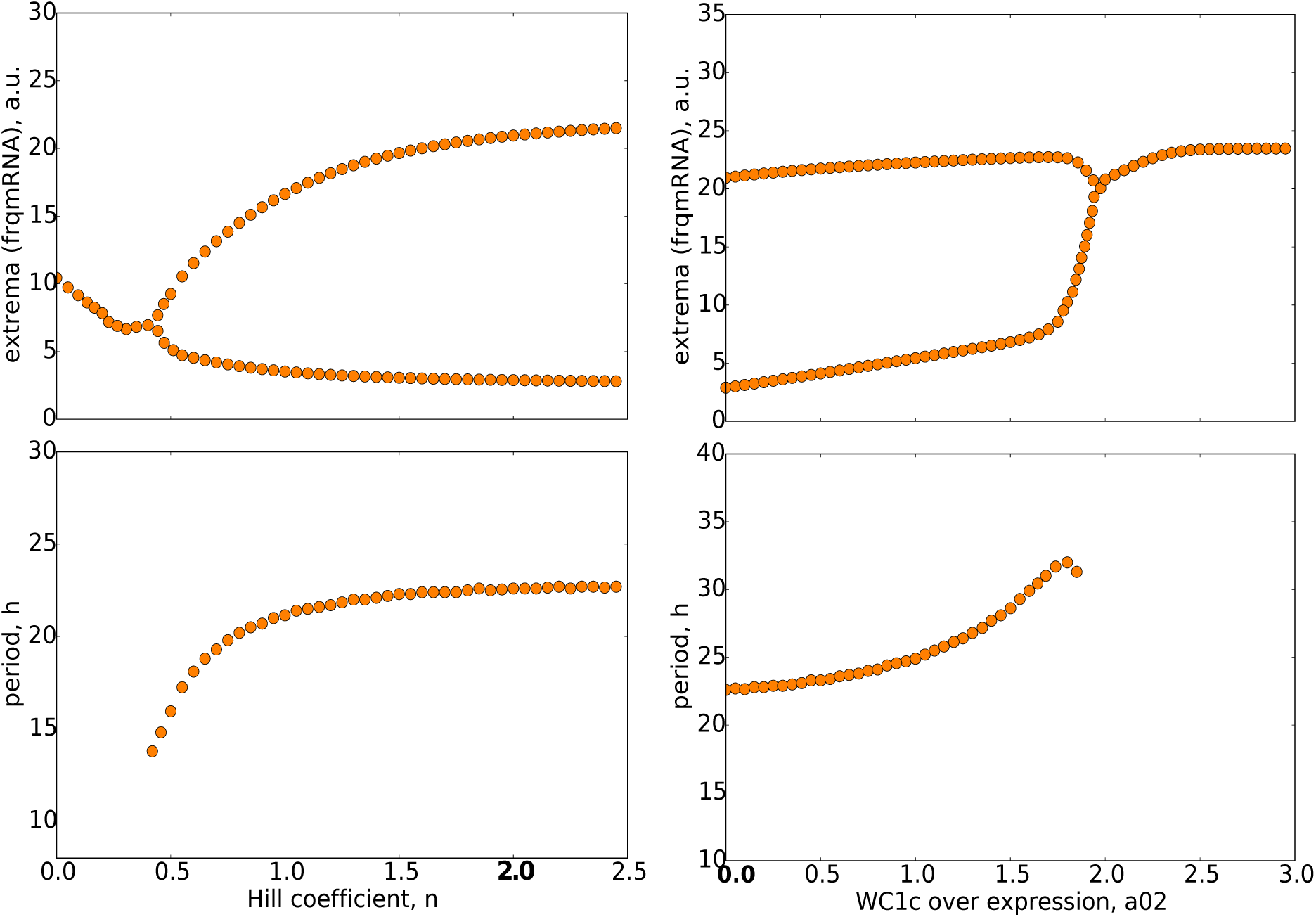
Bifurcation diagrams: The upper graphs show the maxima and minima of oscillations for varying parameters n and a02. The lower graphs depict the corresponding periods. It turns out that oscillations persist for Hill coefficient n between 0.5 to 2.5 whereas overexpression of WC1 can terminate rhythms. Default parameter values are marked by bold numbers.

For 3 parameters we observe hard onsets of oscillations (compare Appendix A3), i.e. the amplitude jumps abruptly. Such bifurcations are characterized by hysteresis and co-existing attractors [39–41](see basins of attractors in Appendix A4). Details of these phenomena are presented in Appendices A3 and A4.

Nonlinearities are necessary to obtain limit cycle oscillations. As discussed above our model exhibits just 3 nonlinear terms. In order to check their relevance we replaced the terms by their mean (termed “clamping” in [42,43]) or by a linearized kinetics. Interestingly, for 2 of the nonlinearities the oscillations persisted. This implies that the dimerization and the positive feedback are not essential to obtain self-sustained oscillations. Thus, the only essential nonlinearity in our model is the Hill function describing frq transcription. Details of the detection of essential nonlinearities are provided in the supplementary information (Appendix A5).

In summary, the developed model exhibits robust self-sustained oscillations with just a single nonlinearity and a fairly small Hill coefficient n=2. Thus we can exploit the model to discuss mutants, to analyze the underlying transcriptional switch, and to explore possible glucose compensation mechanisms.

### 2.4. Our model reproduces clock mutants

*N. crassa* became a model organism in chronobiology since a band mutation in the ras-1 pathway leads to an overt periodic conidiation phenotype. A growth front of rhythmic conidiation and mycelium in so-called race tubes (solid culture) and gene expressions from bioluminescence signals using a frq-promoter luciferase reporter assay help visualizing a period of about 22.5 hours [44–46]. These techniques also allowed the identification of point mutations in the frq gene with altered period lengths. For example, the frq7 mutation stabilizes FRQ leading to a longer period of about 29 hours [47].

Table 1 lists several mutant phenotypes and associated experimental findings regarding protein stability. If we adapt our model rate constants accordingly we can simulate the phenotypes of wildtype (frq+), shorter and longer period mutants (frq1 and frq7) as shown in Appendix A6. Moreover, our model also reproduces the behaviour of a FWD-1 knockout experiment demonstrating that complete turnover of FRQ is not required for the clock to run [48] (see Appendix A6).

**Table 1.**
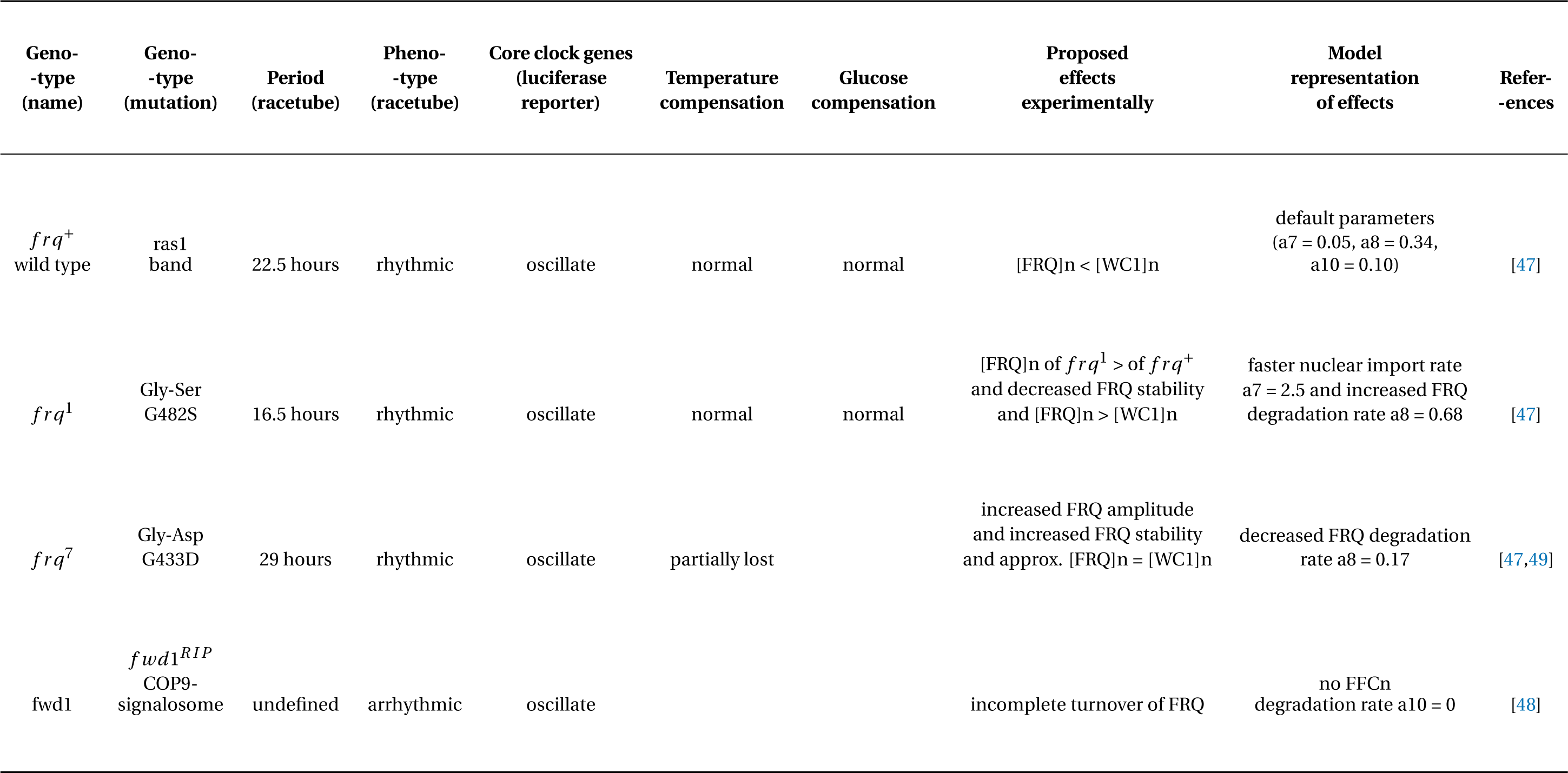
Modeling *Neurospora* clock mutants: A list of selected mutants and their properties (simulations in Appendix A6).

| Geno-<br>-type<br>(name) | Geno-<br>-type<br>(mutation) | Period<br>(racetube) | Pheno-<br>-type<br>(racetube) | Core clock genes<br>(luciferase<br>reporter) | Temperature<br>compensation | Glucose<br>compensation | Proposed<br>effects<br>experimentally | Model<br>representation<br>of effects | Refer-<br>-ences |
| --- | --- | --- | --- | --- | --- | --- | --- | --- | --- |
| *frq*<sup>+</sup><br>wild type | ras1<br>band | 22.5 hours | rhythmic | oscillate | normal | normal | [FRQ]n < [WC1]n | default parameters<br>(a7 = 0.05, a8 = 0.34,<br>a10 = 0.10) | [47] |
| *frq*<sup>1</sup> | Gly-Ser<br>G482S | 16.5 hours | rhythmic | oscillate | normal | normal | [FRQ]n of *frq*<sup>1</sup> > of *frq*<sup>+</sup><br>and decreased FRQ stability<br>and [FRQ]n > [WC1]n | faster nuclear import rate<br>a7 = 2.5 and increased FRQ<br>degradation rate a8 = 0.68 | [47] |
| *frq*<sup>7</sup> | Gly-Asp<br>G433D | 29 hours | rhythmic | oscillate | partially lost |  | increased FRQ amplitude<br>and increased FRQ stability<br>and approx. [FRQ]n = [WC1]n | decreased FRQ degradation<br>rate a8 = 0.17 | [47,49] |
| *fwd1* | *fwd1*<sup>RIP</sup><br>COP9-signalosome | undefined | arrhythmic | oscillate |  |  | incomplete turnover of FRQ | no FFCn<br>degradation rate a10 = 0 | [48] |

### 2.5. An inactivation-switch allows self sustained oscillation

It is widely known that molecular switches support limit cycle oscillations [50–53]. Moreover, the switch-like waveforms in the time series of our model discussed above indicate a switch. However, the biological mechanism of such a switch is not immediately evident.

Multiple mechanisms have been discussed that allow switch-like behaviour. For example, cooperativity [54,55], zero-order ultrasensitivity [56] and multiple phosphorylations [57,58] can lead to quite steep input-output relationships [59]. Furthermore, sequestration [60] and positive feedbacks [61] can generate bistable switches [62–65].

A pure sequestration of the activator WC1n by the FCC complex seems unlikley since the FFCn levels are most of the time lower than the WC1n levels and only a small fraction of WC1n is bound to FFCn (yellow line in Figure 5). Inspection of the rate constants (Appendix A2) reveals the underlying switch mechanism: The degradation rate a18 of the active WC1n is quite low. If the levels of FFCn grow around time 10 in Figure 5, fast binding of FFCn to WC1n (a19 > a20) allows the sudden deactivation of WC1n (a21 > a18). A switch between active transcription to FFC assisted fast inactivation serves as the basic mechanism. Note, that these steps require neither high Hill coefficients nor positive feedbacks as in many other oscillator models [66,67].

**Figure 5.**
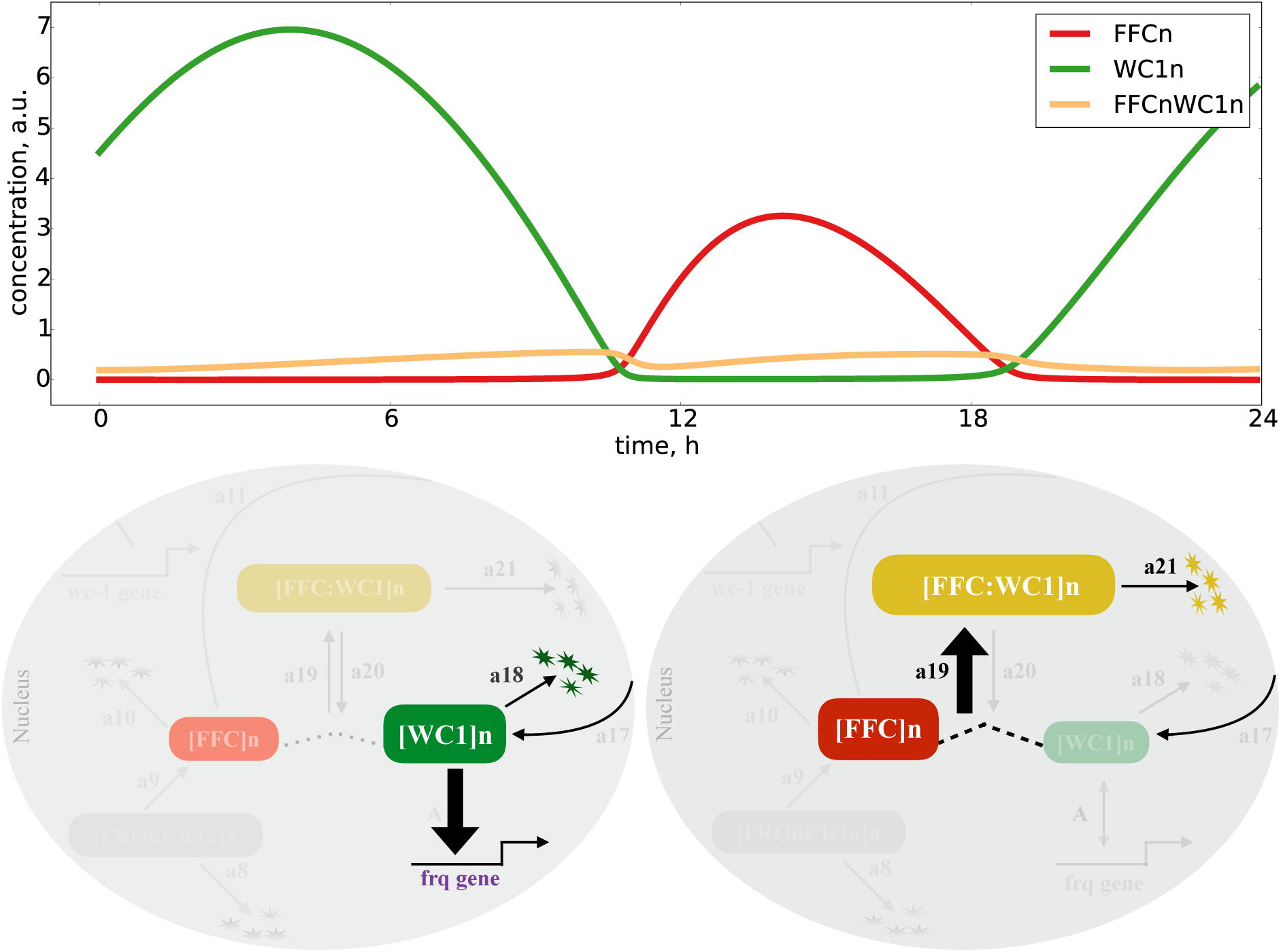
Inactivation switch in *Neurospora* clock model: A switch between WC1n induced transcription and FFC assisted inactivation of WC1n.

### 2.6. Sensitivity analysis points to inherent glucose compensation mechanisms

Circadian clocks, in general, have to be robust against environmental fluctuations. Therefore, they utilize compensation mechanisms against the changes in temperature and nutrients (glucose), where changes in these environmental signals do not affect the overall clock period. There are suggested temperature compensation mechanisms [68,69] and glucose compensation via an auxiliary feedback [70].

Here we study the possibilities for glucose compensation without an extra feedback using our model. We assume that high glucose facilitates transcription and translation of FRQ and WC1. Simulations show that faster transcription and translation rates (a1,a3) of Frequency increase the oscillatory period (compare Figure 6). This is consistent with the data from the mammalian circadian clock [71,72]. With a systematic sensitivity analysis we can search for putative compensation mechanisms. A promising candidate is the helicase FRH since it has a dual role in the network: degradation of nascent mRNAs and formation of FFC complex [27,73].

**Figure 6.**
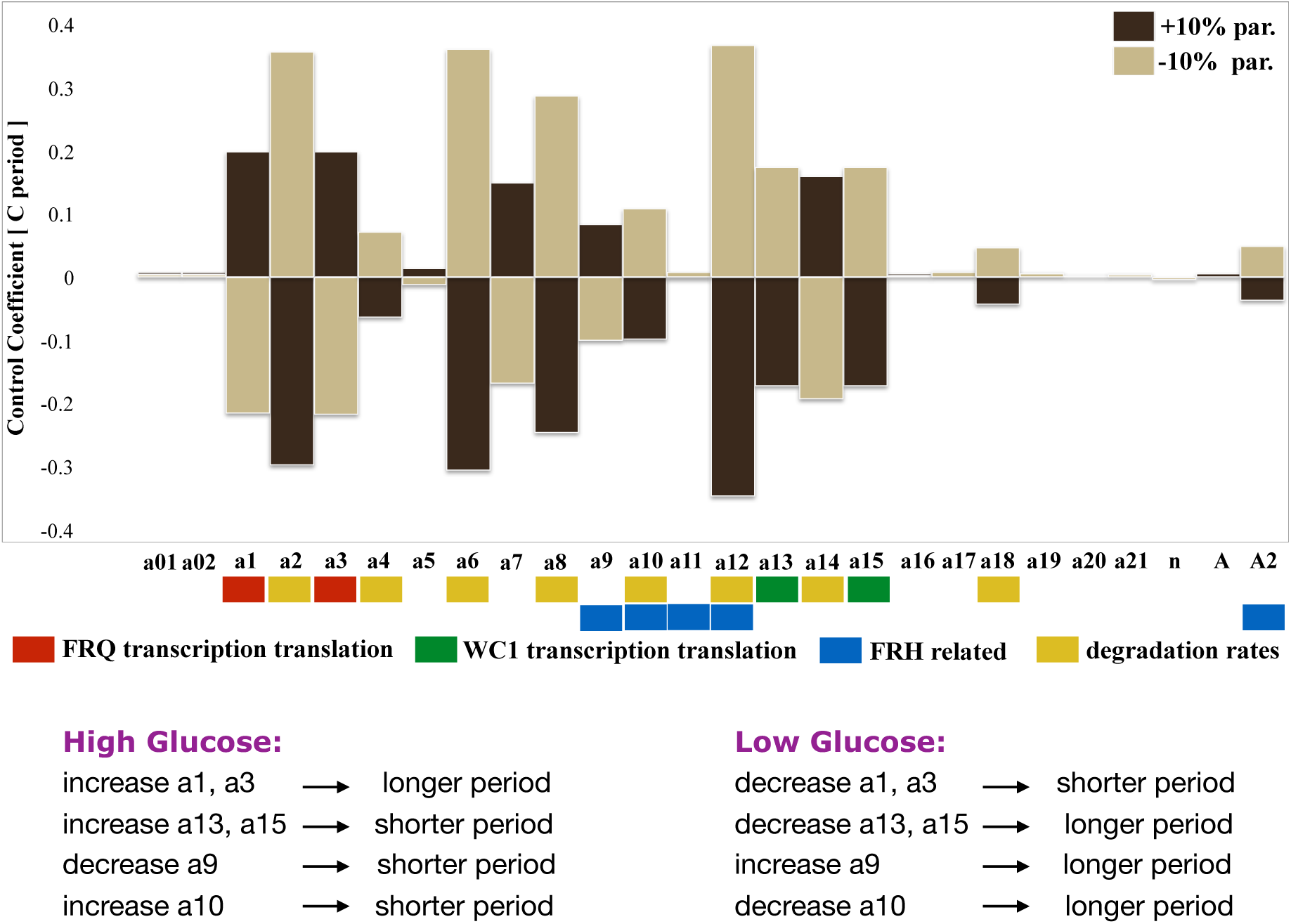
Sensitivity analysis: All parameters were changed by +/− 10 percent and the resulting periods were calculated. Positive and negative period control coefficients indicate period lengthening and shortening respectively.

To quantify period changes for varying the parameters, we calculate the control coefficients quantifying the sensitivity of the system. In Figure 6 we show a comprehensive control analysis of our model along the lines of [37,74,75]. All parameters were changed by +/− 10 percent and the corresponding period changes were extracted from simulations. A positive period control coefficient of 0.5 implies that a 10 percent parameter increase induces a period lengthening by 5 percent, i.e. about 1 hour. Our analysis confirms the well-known feature that faster degradation of Frequency leads to shorter periods [76] as discussed above for the *f rq*^1^ mutant (see the yellow bars associated with parameters a2, a4, a6, a8 and a10).

Now we discuss the possible mechanisms to get compensation of high glucose levels. The period increase via faster Frequency production (a1, a3) could be compensated by faster WC1 production (a13, a15) since the corresponding control coefficients have opposite signs. In order to study the dual role of FRH we marked in Figure 6 the FRH related parameters blue (a9, a10, a11, a12 and A2). It turns out that the dual role of FRH can indeed serve as a compensation mechanism as follows: enhanced transcription due to high glucose sequesters the helicase FRH. This leads to a slower assembly of FFC (smaller a9 value) and less stabilization of FFC (larger a10 value). According to Figure 6 these effects can compensate the period increase due to an increased Frequency production. Interestingly, the glucose compensation through the dual role of FRH is lost if we cancel the positive feedback of FFCc on WC1c translation via clamping (Appendix A7).

## 3. Discussion

Understanding circadian clocks quantitatively helps to explore its functional significance in different organisms. In mammals, a functional clock can help to prevent diseases such as cancer, obesity, and depression [4]. To reveal the basic of mechanisms of the underlying gene-regulatory network (see Figure 1) *Neurospora crassa* has been established as a useful model organism [2].

Mathematical modeling complements experimental studies and can help to uncover the underlying design principles [12]. A transcriptional-translational feedback loop (TTFL) serves as the generator of self-sustained oscillations with a period of about a day but quantitative details of these delayed negative feedbacks [77] and supporting molecular switches [78] are debated.

The required delay of at least 6 hours [79] is associated with gene expression, phosphorylation, nuclear translocation, complex formations and epigenetic regulations [80,81]. In addition to appropriate delays limit cycle oscillations require nonlinearities such as molecular switches [37] as studied extensively in cell proliferation and differentiation [50,62].

To explore the temporal dynamics of the *Neurospora* clock, we developed a mathematical model (Figure 2) that reproduces basic features of wild type rhythms and selected mutants (Table 1). Simulations reveal spike-like waveforms and a temporal transcriptional switch (Figure 3). Systematic clamping and linearization reveals that there is just one essential nonlinearity with a Hill coefficient of n=2. This raises the question how the ON/OFF switch of the Frequency gene is generated without explicit ultrasensitivity or bistability.

As illustrated in Figure 5 we find sharp transitions between active transcription with slow turnover of WC1 and fast FFC assisted inactivation of WC1. Such a protein inactivation switch has also been discussed in recent studies [82–85].

Our model can also be exploited to study compensations of environmental fluctuations. It has been found experimentally that the *Neurospora* clock keeps its period almost constant at varying temperatures and glucose concentrations [68,69]. In a previous study an auxiliary transcriptional feedback loop has been postulated for glucose compensation [70].

Sensitivity analysis of our model (Figure 6) suggests in-built mechanisms without additional feedbacks. High levels of glucose enhance gene expression. Our model indicates that higher levels of core clock genes increase the period as found experimentally in *Neurospora* [6,14] and in mammals [71]. Interestingly, enhanced WC1 levels decrease the period (compare Figure 6). This counter-regulation constitutes a first putative compensation mechanism.

Another glucose compensation mechanism is based on the dual role of FRH. This protein serves as an RNA helicase and as a stabilizer of the FFC complex [27,73]. Since at high glucose conditions many more mRNAs are transcribed FRH is partially sequestered. This implies that the FFC complex is destabilized leading to a shortening of the period. This effect can compensate the increased period induced by enhanced transcription. Interestingly this compensation is diminished if we turn off the positive feedback via WC1 translation [25]. This finding points to a possible role of this additional feedback regulation. Note, that this feedback was quite weak in the original model [32]. We adjusted the parameter A2 to activate in our model this experimentally verified positive feedback [25].

In summary, our proposed mathematical model with 10 dynamic variables and 26 kinetic parameters reproduces many experimentally known features. A comprehensive analysis of the model reveals some unexpected results. It turns out that there is just a single nonlinearity essential for self-sustained oscillations. The switch-like behavior of the model arises from a novel inactivation switch based of WC1 inactivation assisted by the FFC complex. This finding suggests that forthcoming experiments should be oriented towards the detailed mechanisms of WC1-FFC interactions.

Another result concerns the compensation of environmental perturbations. Sensitivity analysis of our model suggests putative in-built mechanisms to achieve glucose compensation. First expression of the WC1 gene could counteract the effects of Frequency expression on the period. Another mechanism points to possible roles of FRH and the positive feedback. Sequestration of FRH via increasing mRNA transcription leads to a destabilization of the FFC complex which can compensate period changes in the presence of the positive feedback.

Even though our model is not quantitative in any detail the comparison of simulations with experimental data can stimulate future investigations. The combination of a delayed negative feedback with an inactivation switch is a robust design principle that could be relevant also in other oscillatory systems such as cell cycle dynamics [86], redox oscillations [87] or somite formation [88].

## 4. Materials and Methods

All the simulations have been performed using codes on a Spyder Python 3.4 platform. Ordinary differential equations have been written into codes using SciPy odeint library. Matplotlib library has been used to generate figures. XPP-Auto has been used to reproduce the bifurcation diagrams generated by Python. Codes are available on request.

## Author Contributions

conceptualization by M.B. and H.P.; methodology by A.U. and H.P.; software by A.U.; validation by A.U. and H.P.; formal analysis by A.U.; investigation, A.U. and H.P.; resources by H.P.; data curation by M.B.; writing—original draft preparation by A.U. and H.P.; writing—review and editing, A.U., H.P. and M.B.; visualization by A.U. and H.P.; supervision by M.B. and H.P.; project administration by M.B. and H.P.; funding acquisition by M.B. and H.P.

## Acknowledgments

Authors are thankful to J Patrick Pett, Christoph Schmal, Daniela Marzoll, Axel Diernfellner, Norbert Gyongyosi, Marta del Olmo, Philipp Burt and Katharina Baum for their fruitful discussions.

## Abbreviations

The following abbreviations are used in this manuscript:

*N. crassa*: *Neurospora crassa*
FRQ: Frequency protein
WCC: White Collar Complex
CK1a: Casein Kinase 1a
FRH: Frequency-interacting Helicase
FFC: Frequency FRH CK1a complex
FWD1: F-box/WD-40 repeat-containing protein
a.u.: arbitrary units
ODEs: Ordinary Differential Equations

## Appendix A

**Figure A1.**
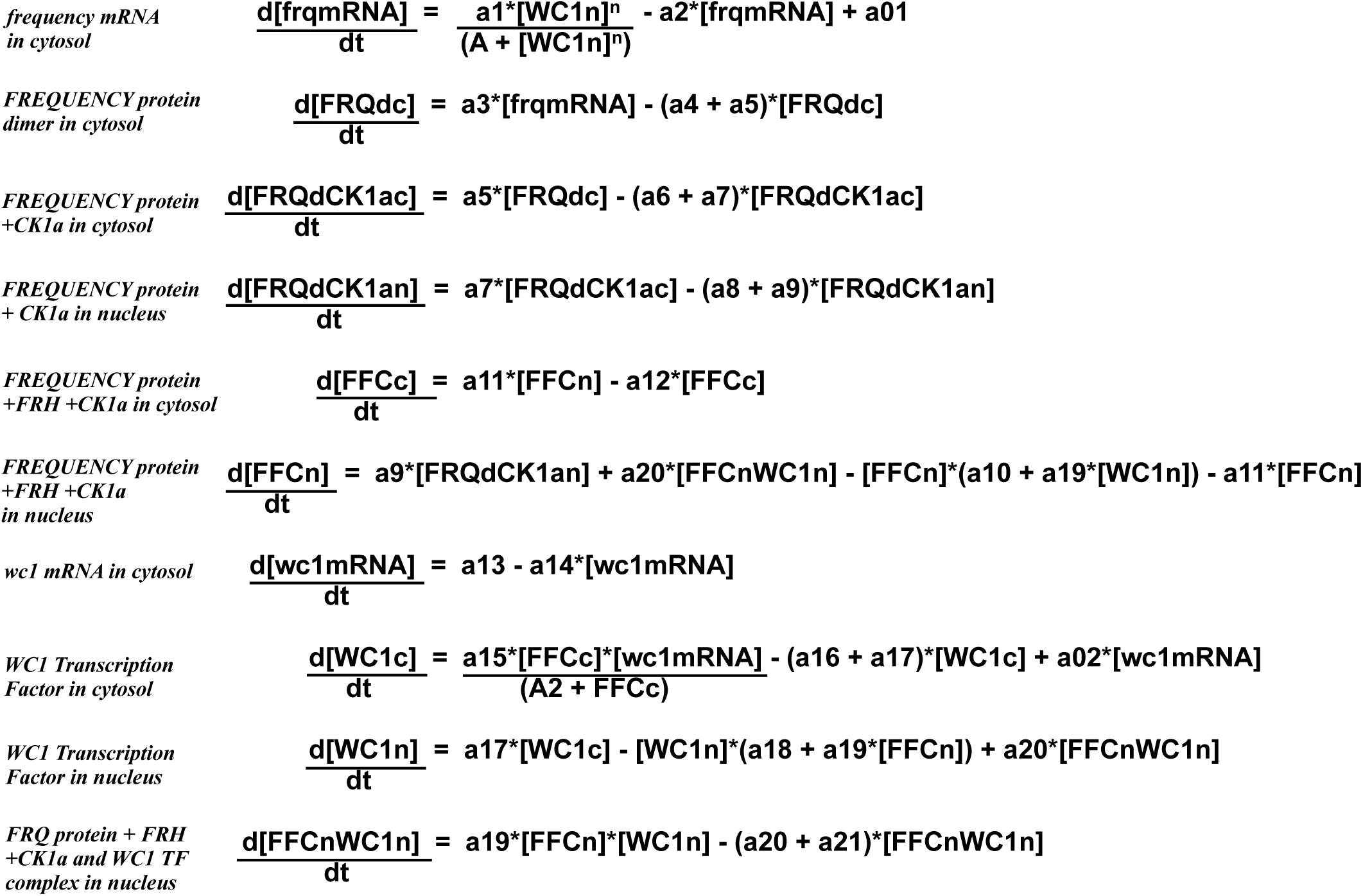
Ordinary differential equations of model: A 10 variables and 26 parameters model.

**Figure A2.**
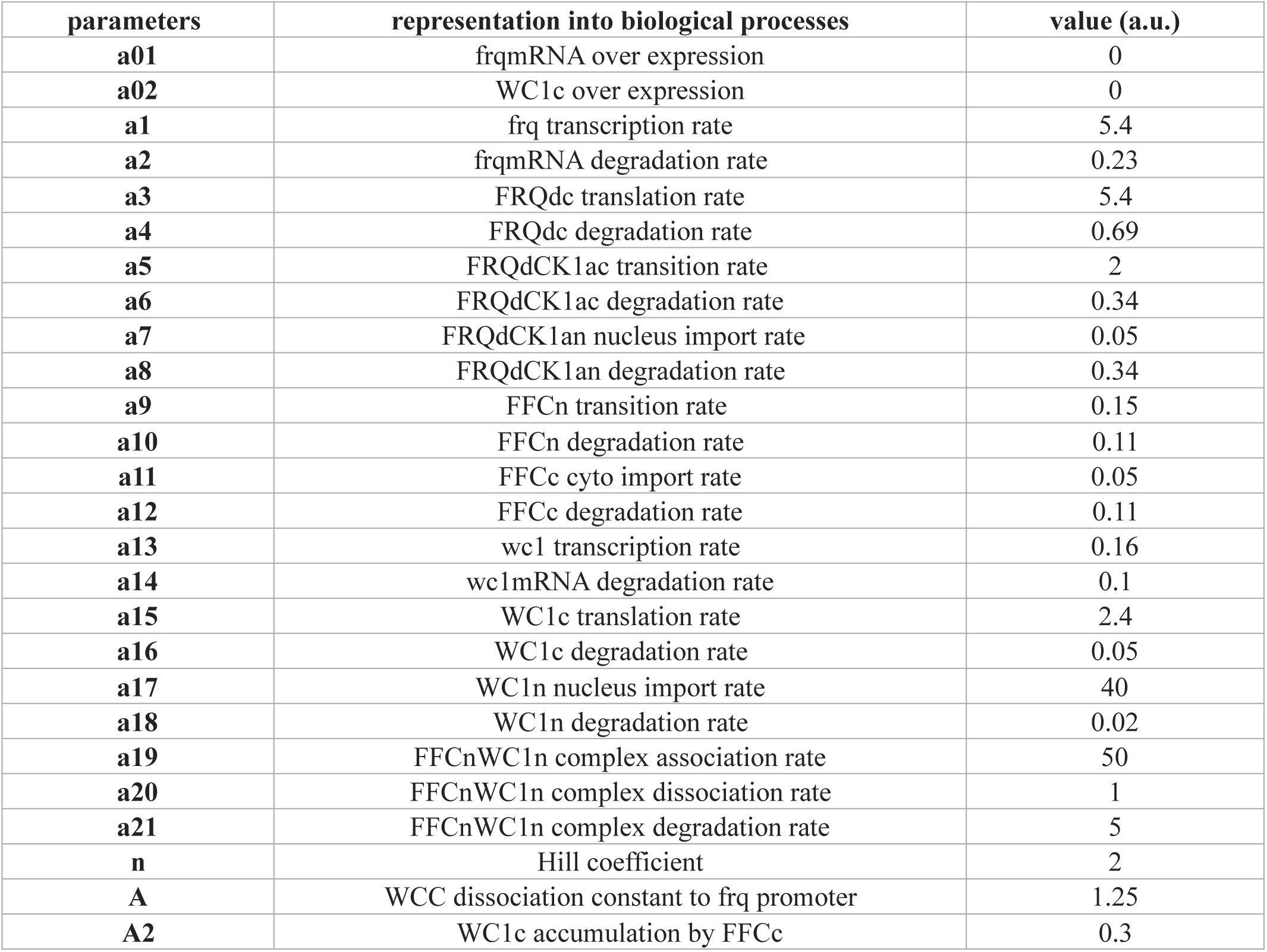
Kinetic parameters: A list of rate constants and their associated biological processes.

**Figure A3.**
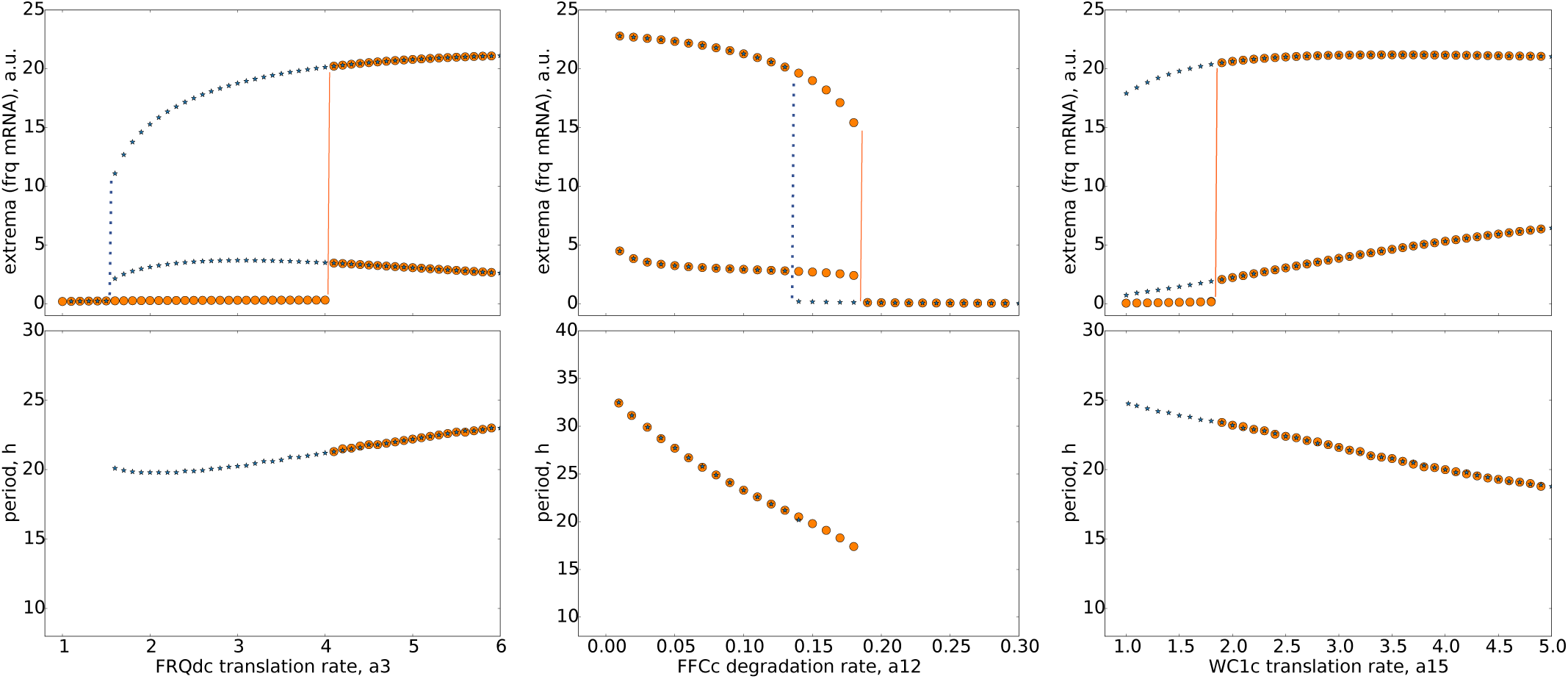
Co-existence of attractors: Bifurcation analysis of three parameters shows co-existence of limit cycle and steasy state while parameter is slowly varied. The values for increasing parameters are marked by orange circles. Decreasing parameters correspond to blue stars. Red lines mark jumps between steady states (low frq values) to limit cycle oscillations. The associated periods are given in the lower graphs.

**Figure A4.**
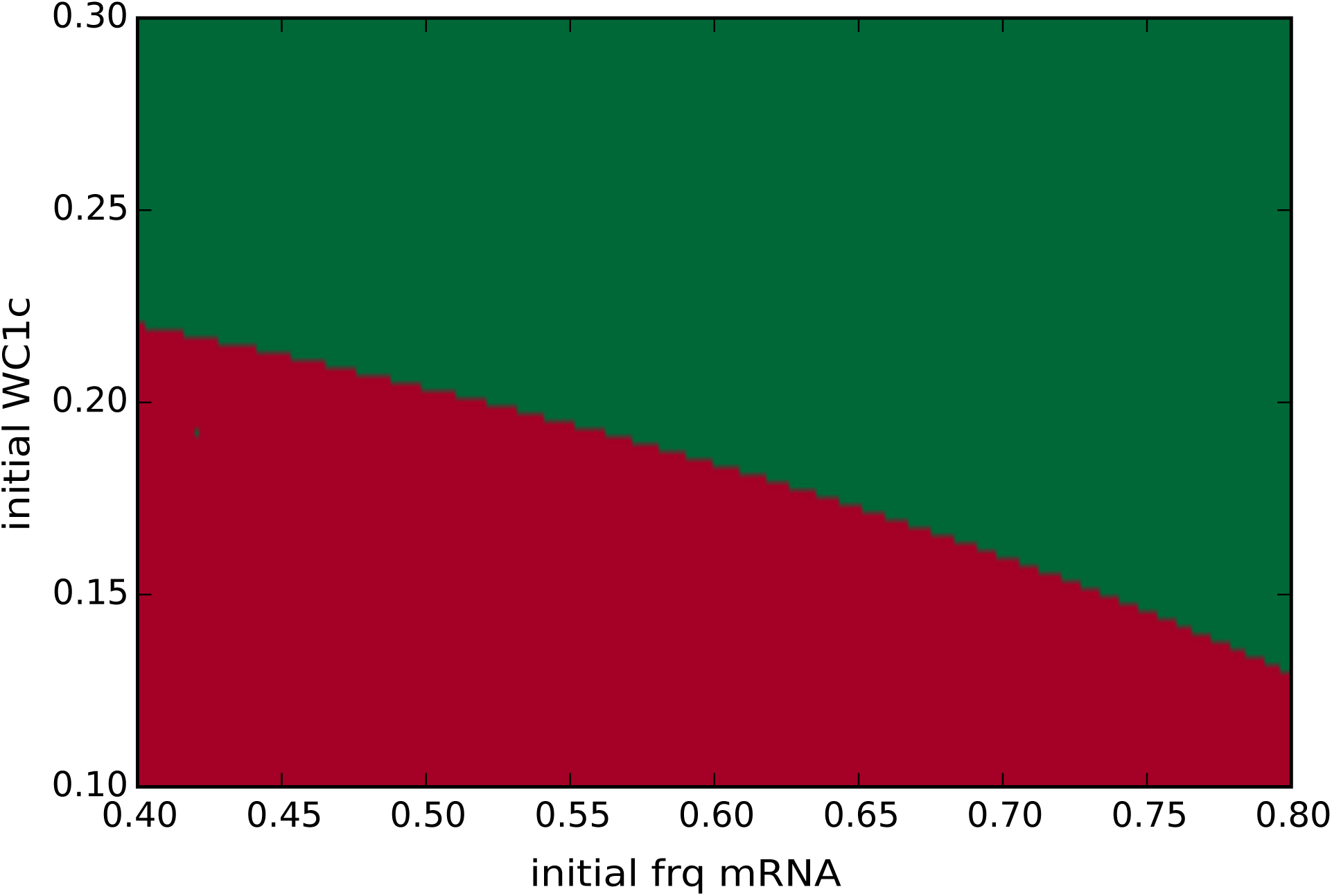
Basins of attractors: Coexisting attractors shown in ‘Appendix A3’ imply distinct basis of attraction (here a3 = 3). Small initial values of frq and wc1 (red area) approach the steady state whereas initial conditions in green approach the limit cycle. The initial conditions of the rest of the variables were zero.

**Figure A5.**
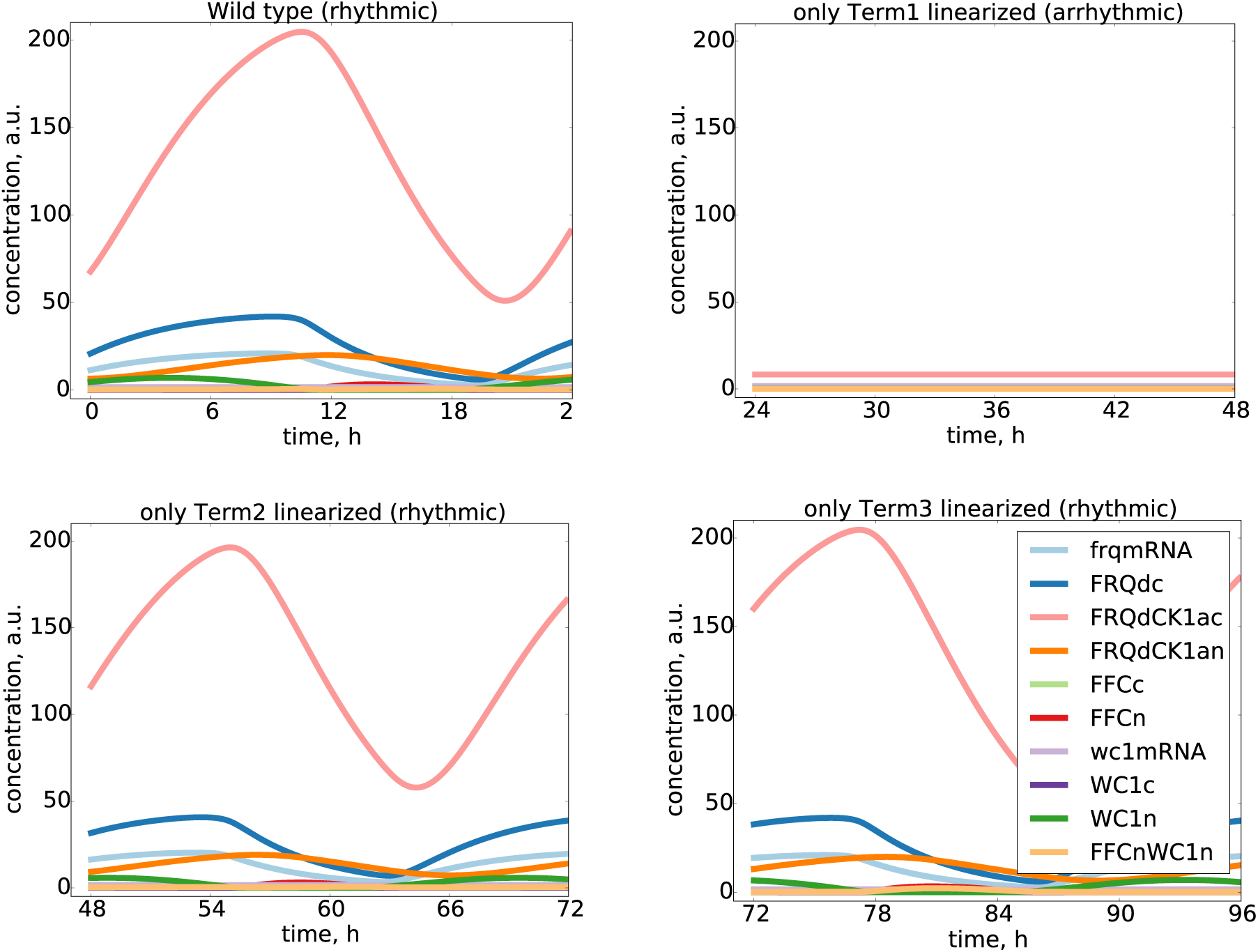
Essential and non-essential nonlinearities in the model: We linearized all the 3 nonlinear terms and find that the Hill function of frq production (Term1) is essential whereas the positive feedback nonlinearity (Term2) and the bilinear dimerization (Term3) are not essential since linearizations kept the rhythmicity.

**Figure A6.**
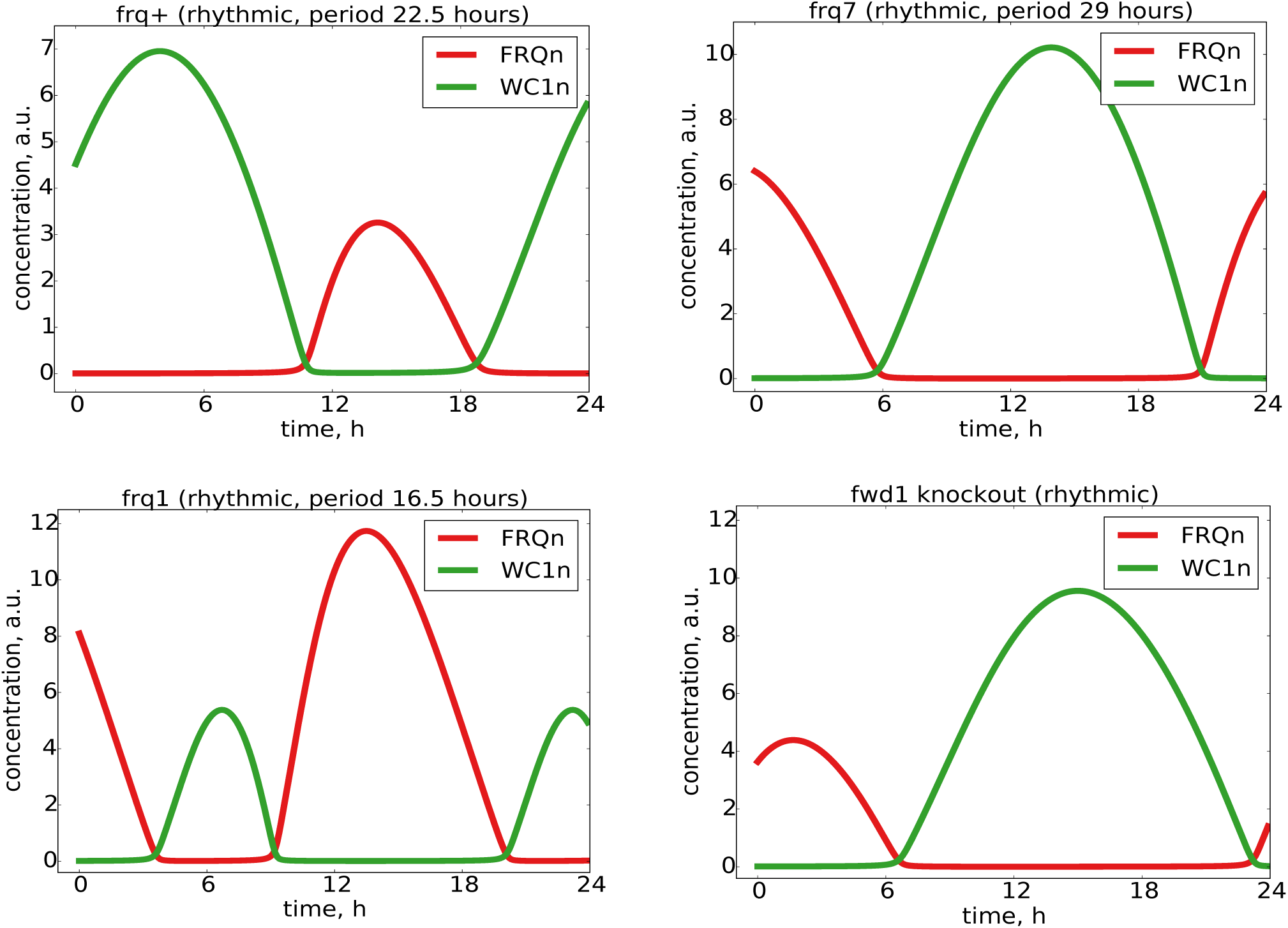
Mutants simulations: Several experimentally observed mutants are also reproduced by our model. The corresponding modified parameters are given in Table 1.

**Figure A7.**
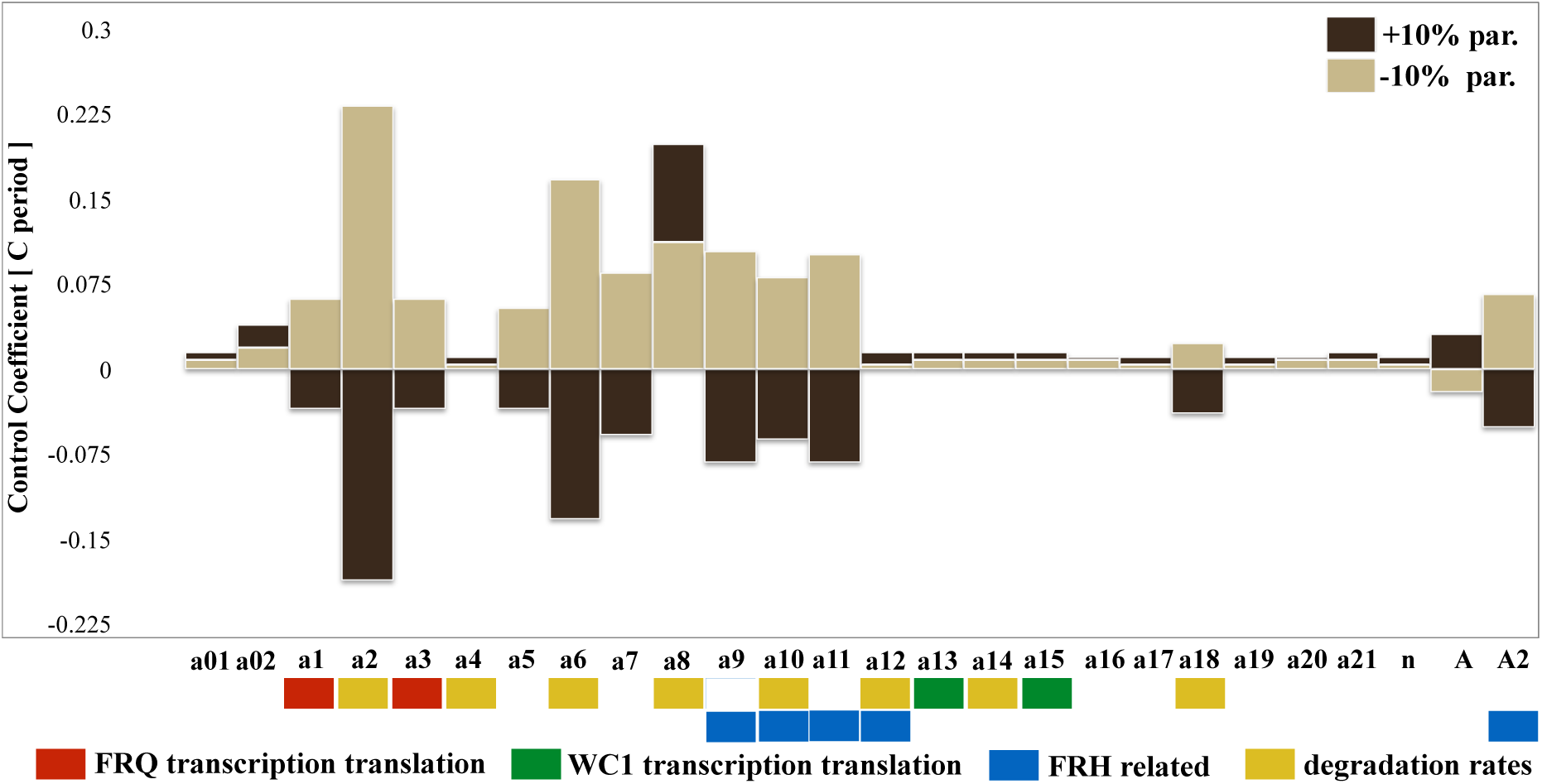
Sensitivity analysis after clamping the positive feedback: All parameters were changed by +/− 10 percent and the resulting periods were calculated. Positive and negative period control coefficients indicate period lengthening and shortening respectively (compare Figure 6).

